# Decoding functional regulatory maps via genomic evolutionary footprints in 63 green plants

**DOI:** 10.1101/484493

**Authors:** Feng Tian, De-Chang Yang, Yu-Qi Meng, Jinpu Jin, Ge Gao

## Abstract

Systematic identification of functional transcriptional regulatory interactions is essential for understanding regulatory systems. Here, we firstly established genome-wide conservation landscapes for 63 green plants of seven lineages and then developed an algorithm FunTFBS to screen for functional regulatory elements and interactions by coupling base-varied binding affinities of transcription factors with the evolutionary footprints on their binding sites. Using the FunTFBS and the conservation landscapes, we further identified over two million functional interactions for 21,346 TFs, charting functional regulatory maps of these 63 plants. Our work provides plant community with valuable resources to decode plant transcriptional regulatory system and genome sequences.

## Background

Transcription factors (TFs) control gene expression by binding to specific cis-elements, which play essential roles in plant development and stress responses. A systematic identification of the functional transcriptional regulatory interactions would greatly facilitate further mechanism investigation [1, 2]. With the binding motifs of genome-wide TFs determined by experiments in plants [3, 4] and *in silico* mapped into 156 plants [5], directly scanning the TF binding motifs in the promoters of putative target genes becomes a promising option to chart plant transcriptional regulatory maps. As prediction from directly scanning yields a rather high false positive rate, additional data like DNase-seq footprints [6, 7] and conserved elements [8–10] have been incorporated to screen for functional TF binding sites (TFBSs). However, these data are only available in a few model plants [7, 11] and conserved element-based methods are still confounded by evolutionary constraints from other functional elements besides of TF binding [12], hindering this filed from systematically charting the transcriptional regulatory maps across the plant kingdom.

A comparison of multiple related genomes with substantial divergence is widely used to detect evolutionary constraints and further identify the functional elements [11, 13, 14]. The availability of over one hundred of plant genomes provides a unique opportunity to calculate genome-wide evolutionary footprints and further to chart functional plant regulatory maps. In this study, we established the first genome-wide conservation landscapes for 63 green plants and developed an algorithm by coupling base-varied binding affinities of TFs with the evolutionary footprints on their binding sites to screen functional transcription regulatory elements and to infer interactions, charting the regulatory maps of these 63 plants.

## Results and Discussion

To detect evolutionary constraints, we first chose 63 representative species of seven lineages (Fig. 1a), with the divergence time from 37 million years ago (MYA) to 106 MYA (Fig. 1b), covering main lineages of green plants. Then, we performed all 322 intragroup pairwise genome alignments and then assembled them into 63 multiple genome alignments by using each species as a reference genome (see Additional file 1: Supplementary Methods for more details). Finally, we identified over 67 million conserved elements and calculated the basewise conservation scores for over 22 billion base pairs, covering approximately 66% of genome sequences and establishing the first conservation landscapes covering the main lineages of green plants (Fig. 1b; Additional file 1: Tables S1 and S2).

**Fig. 1.**
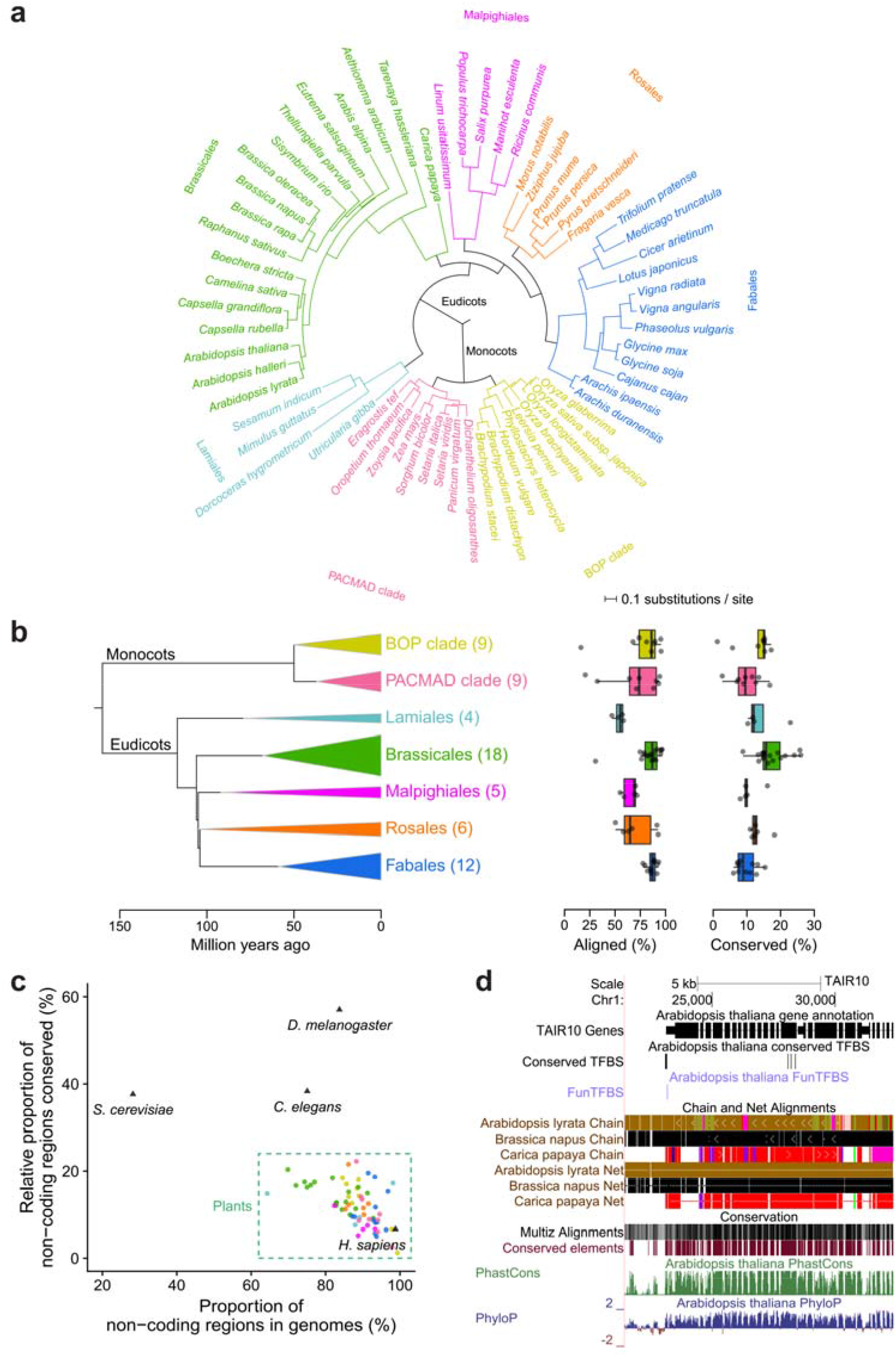
Genome-wide conservation landscapes of 63 green plants. **a**, Phylogenetic tree for 63 species of seven lineages, where the branch length within each group is shown as the substitution number per site at four-fold degenerate sites. **b**, The summary of genomes aligned and conserved for each group. Left: The phylogenetic tree shows the evolutionary relationship among groups. The width of the triangle corresponds to the maximum divergence time of involved species, and the height corresponds to the species number (as shown in brackets) in each group. Middle and Right: The box plots show the percentage of genomes aligned (Middle) and conserved (Right) for each group. **c**, The proportion of non-coding regions in the genomes and their relative conservation ratio (proportion of non-coding regions conserved / proportion of coding regions conserved) for plants (63 species), animals (*H. sapiens, D. melanogaster* and *C. elegans*) and yeasts (*S. cerevisiae*). Each data point represents a species and its color represents the group. **d**, The screenshot of genome alignment and conservation resource in the genome browser.

For these seven groups, approximately 54-87% of genomes are aligned together, and at least 10-17% of the genomes are under evolutionary constraint (Fig. 1b; Additional file 1: Tables S1 and S2). Compared with a previous study in *A. thaliana* [11] which used nine species in *Brassicales* to calculate conservation scores, our work, which embraced more representative species (18 *vs* 9), aligned 20.49% (18.13 Mb) more genome sequences and detected 100.96% (4.84 Mb) more conserved non-coding regions with a higher accuracy (Additional file 1: Figure S1). Compared with other kingdoms, plants present a higher conserved ratio in the noncoding regions than these of humans but less than those organisms with lower proportion of non-coding sequences in genomes, such as fruit flies, worms, and yeast (Fig. 1c). Lineages suffering more rounds of whole genome duplication (e.g., *Fabales*) or sudden genome expansion (e.g., *PACMAD clade*) show lower conservation ratios in their genomes, likely due to genomic degeneration after whole genome duplication or a lack of homologs in close species after sudden expansion (e.g., *Zea mays* [15]) (Fig. 1b, c; Additional file 1: Table S1). Of note, compared to the unannotated nonconserved regions, the unannotated conserved regions in *A. thaliana* genome present a larger proportion covered by transcriptional signals (27% *vs* 10%) and a higher expression level (Additional file 1: Figure S2), suggesting that many genes or functional elements still remain to be illustrated even in this most well-annotated plant genome. Conservation landscapes in humans and fruit flies have been widely used to illustrate their genome sequences [13, 14], thus our conservation landscapes in 63 plants provide the community a unique chance to illustrate plant genomes. For users to conveniently access the conservation data, we have set up a genome browser to visualize each species (Fig. 1d) [16].

The establishment of conservation landscapes in the main lineage of green plants paves the way for a systematic identification of functional TFBSs. However, other functional elements besides of TFBSs (such as noncoding RNAs and stem regions in RNA structures) may also result in a conservation at the gene promoters [12] (Additional file 1: Figure S3), confounding algorithms that depend on conserved elements to screen for functional interactions. We speculate that the base-varied binding affinity (base frequencies in the binding motifs) of the TF binding motifs could yield a consistent base-varied evolutionary constraint on the functional TFBSs (Fig. 2a). To determine whether this feature could distinguish functional TFBSs from nonfunctional ones, we firstly featured out an evaluation dataset by classifying the TFBSs that were identified from 124 ChIP-seq experiments for 21 TFs [17] into three classes: “Less reliable”, “Highly reliable” and “Functional”. The “Less reliable” and the “Highly reliable” TFBSs represent the TFBSs with low and high consistency among replicates, respectively, and the “Functional” ones are the “Highly reliable” TFBSs that are further supported by expression data (Additional file 1: Supplementary Methods). A method with a higher screening efficiency would result in a lower percentage of TFBSs supported by “Less reliable” TFBSs but a higher percentage of TFBSs supported by “Highly reliable” and “Functional” TFBSs. Compared with the TFBSs whose conservation scores are inconsistent with their binding affinity, the consistent TFBSs are depleted in the “Less reliable” TFBSs and enriched in the “Highly reliable” TFBSs, particularly the “Functional” ones (Fig. 2b), suggesting that this feature does work to screen for functional TFBSs.

**Fig. 2.**
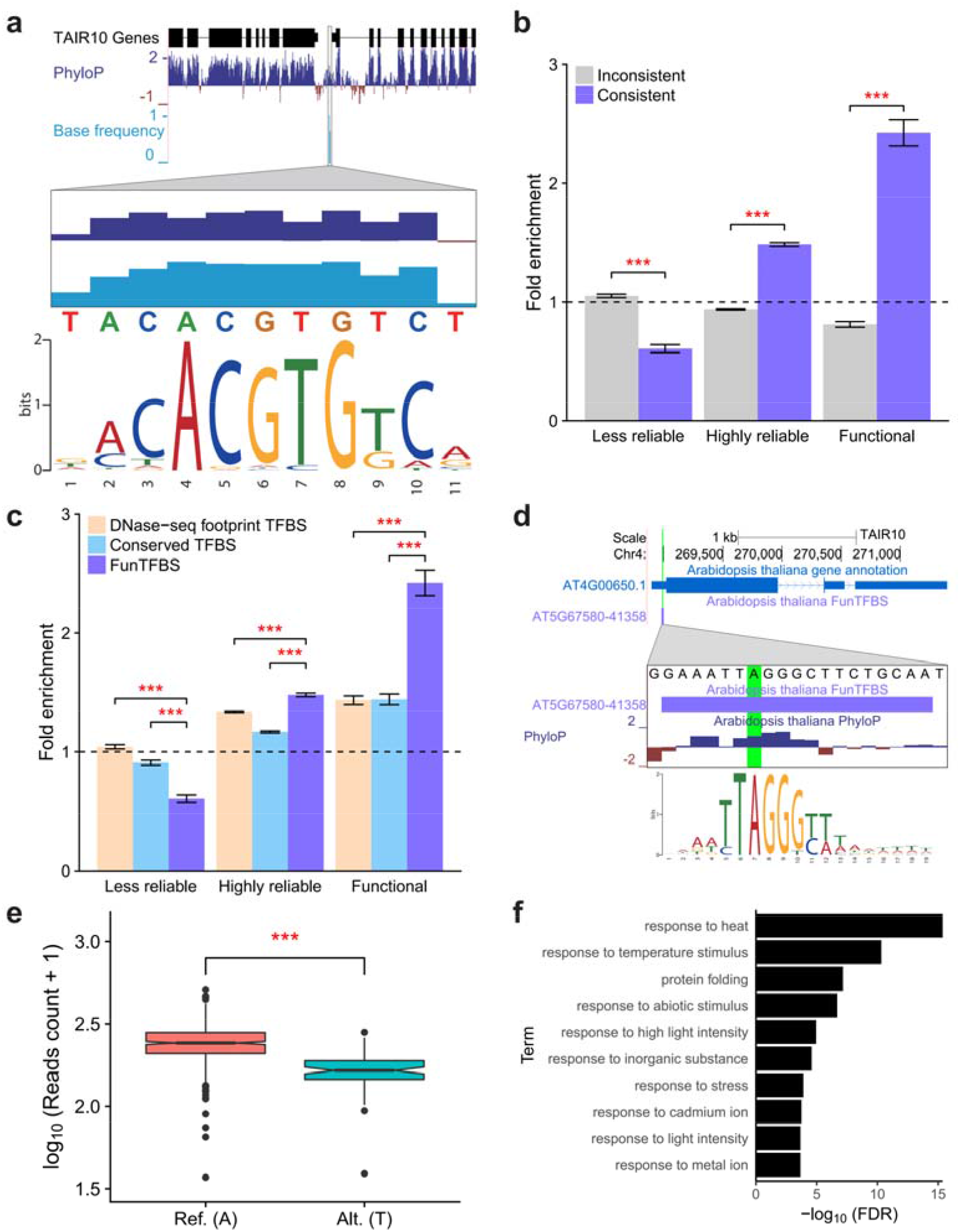
Screening for functional regulatory interactions by coupling the base-varied binding affinities of transcription factors (TFs) and consistent evolutionary constraints on their binding sites (TFBSs). **a**, An example illustrates the consistency between the based-varied binding affinities (base frequency in the binding motifs) of the TFs and the evolutionary constraints on a functional TFBS. **b and c**, Enrichment analysis of the TFBSs showing consistency or inconsistency between conservation scores and base frequencies in motifs (**b**) and the TFBSs screened by DNase-seq footprints (DNase-seq footprint TFBS), conserved elements (Conserved TFBS), and evolutionary footprints (FunTFBS) (**c**) in “Less reliable”, “Highly reliable”, and “Functional” TFBSs of the evaluation dataset. The percentage of the TFBSs, directly scanned by motifs, supported by each class of evaluation dataset is used as background. (Error bar indicates the standard deviation of the average fold change from 1000 subsampling; p-value was calculated based on the results of the 1000 subsampling, *** p-value < 0.001.) **d and e**, An eQTL (A to T substitution, highlighted in green) located on a TFBS of AT5G67580 predicted by FunTFBS (**d**) and the significant difference in the expression level of its target gene AT4G00650 (**e**) (Wilcoxon rank sum test, *** p-value < 0.001.) **f**, Enriched GO terms for the target genes of AT3G22830 inferred by FunTFBS.

Employing this feature, we developed an algorithm called FunTFBS to screen for functional TFBSs and to infer the functional regulatory interactions (Additional file 1: Figure S4 and Supplementary Methods). To determine whether our algorithm showed a higher accuracy for functional TFBSs than the other motif-based methods, we firstly compared FunTFBS with the existing DNase-seq footprint-based and conserved element-based methods using the evaluation dataset mentioned above. The result showed that our algorithm presented a 42% and 33% decrease in the percentage of screened TFBSs that were supported by “Less reliable” TFBSs, but it presented a 68% and 67% increase in the percentage of screened TFBSs that were supported by “Functional” TFBSs compared to the DNase-seq footprint-based and conserved element-based methods, respectively, suggesting that our algorithm could more efficiently screen for functional TFBSs (Fig. 2c).

We then assessed the accuracy of our algorithm in inferring transcriptional regulatory interactions based on experimentally validated interactions from *Arabidopsis* transcriptional regulatory map (ATRM) [2]. Our algorithm showed a 95-146% increase in the percentage of edges that were supported by the functional regulatory interactions in ATRM than the DNase-seq footprint-based and conserved element-based methods (Additional file 1: Figure S5), indicating the superiority of FunTFBS in inferring functional regulatory interactions. We further assessed the performance of our algorithm based on another two indexes: the percentage of regulatory pairs that coexist in the same biological process and the percentage of regulatory pairs that are highly correlated in expression [18], where higher numbers in the two indexes represent higher-quality interactions. Our algorithm showed the highest percentage of TFs and their targets coexisting in the two indexes (20-22% and 20-39% increases than the other two methods in the two indexes, respectively) (Additional file 1: Figures S6 and S7), further confirming the superiority of FunTFBS in screening for functional regulatory interactions.

After confirming the superiority of FunTFBS in screening for functional regulatory interactions, we employed this method with integrated genomic TF binding motifs in these 63 plants [5]. Finally, we identified 21,997,501 functional TFBSs and further inferred 2,196,397 regulatory interactions for 21,346 TFs (Additional file 1: Table S3), charting the functional regulatory maps for the main lineages of green plants.

Our identified functional TFBSs are significantly enriched in expression quantitative trait loci (eQTLs) (Additional file 1: Figure S8), offering a unique chance to unveil the molecular mechanisms that underlie genetic variation and gene expression alternation. For example, according to the 1,203 transcriptomes from the 1001 *Arabidopsis* genome project [19], one substitution (Chr4:268990 A>T) is associated with lower expression of AT4G00650, a major source of variation in flowering time. Through browsing our functional TFBSs, we found a TF (AT5G67580) that could bind to that position, and an A to T substitution would definitely weaken its binding (Fig. 2d, e), shedding lights on the putative molecular mechanism. Moreover, the functional regulations also provide informative clues for the function of the TFs. For example, the target genes of the transcription factor AT3G22830 were found to be enriched in “response to heat” (Fig. 2f), a biological process which corresponds well to its reported heat stress response function [20].

Finally, the conservation landscapes, the FunTFBS tool and the screened functional TFBSs and regulatory interactions were integrated into the PlantRegMap [5, 21] (More details see Additional file 1: Table S4), a popular plant transcriptional regulatory data and analysis platform, to better serve the community.

## Conclusions

In this study, we established the first genome-wide, basewise conservation landscapes of 63 green plants in seven lineages. We further developed a novel algorithm by coupling the base-varied binding affinities of the TFs and the evolutionary footprints on their binding sites to screen for functional TFBSs. Using this method, we systematically screened for the functional TFBSs and regulatory interactions in 63 plants, charting functional regulatory maps for the main branches of higher plants. We believe these resources will advance the understanding of plant transcriptional regulatory systems and the further illustration of plant genome sequences.

## Supporting information

## List of Abbreviations

ATRM: : *Arabidopsis* transcriptional regulatory map
eQTLs: : Expression quantitative trait loci
MYA: : Million year ago
TF: : Transcription factor
TFBS: : Transcription factor binding site

## Additional files

Additional file 1: Supplementary Methods, Figures S1-S8, and Tables S1-S4.

## Acknowledgments

Part of the analyses were performed on the Computing Platform of the Center for Life Sciences of Peking University, and we thank Dr. Fangjin Chen and Ting Fang for their helps.

## Availability of data and materials

The conservation landscapes and the regulatory maps for the 63 plant species are available at PlantRegMap (http://plantregmap.cbi.pku.edu.cn/) [5, 21]. The genome browser for data visualization is available at <u>http://plantregmap.cbi.pku.edu.cn/cis-map.php</u> [16]. The FunTFBS tool and the evaluation dataset are available at <u>http://plantregmap.cbi.pku.edu.cn/funtfbs.php</u> [22]. See Additional file 1: Table S4 for more details about the resource accessibility.

## Authors’ Contributions

GG and JJ conceived and designed the research. FT performed all the data analysis and tool development. DCY set up the website for data visualization and user interaction. YQM and DCY set up the genome browser for cis-Map. FT, JJ and GG wrote the manuscript with contribution from all the authors.

## Ethics approval and consent to participate

Not applicable

## Consent for publication

Not applicable

