## Supplementary material for "Decoding functional regulatory maps via genomic evolutionary footprints in 63 green plants"

**Supplementary Methods**

**The workflow for establishing conservation landscapes across green plants**

Genome sequences and gene annotation of 156 plant species were downloaded from PlantTFDB 4.0 (see http://planttfdb.cbi.pku.edu.cn/help_datasrc.php for more details about the data source) [1] and then the species were grouped according to their taxonomy. The groups including no less than four species that presented high-quality genome assembly (63 species in seven groups, Fig. 1a) were kept for further conservation analysis. For each species, RepeatMasker (v4.0.6 with RepBase version ‘20150807’) [2] was employed to mask the transposons and low complexity sequences within the genome. Then, pairwise alignments were performed inside each group using LASTZ (v1.03.73) [3] with the parameters “--inner=2000 --xdrop=9400 --gappedthresh=3000 --hspthresh=2200” referred from a previous study [4]. Pairwise alignment blocks were further chained and netted by axtChain [5] and chainNet [5] to produce single coverage alignment for each target genome. Multiple alignments were assembled using Multiz (v11.2) [6] based on their phylogenetic trees. Conservation scores (PhastCons scores and PhyloP scores) and conserved elements were calculated using RPHAST (v1.6) [7]. For each species, the parameters were tuned until the percentage of conserved elements in the aligned CDS regions was near 65% according to the manual (http://compgen.cshl.edu/phast/phastCons-HOWTO.html). The conserved elements and PhastCons scores were calculated with the “viterbi” mode, and the PhyloP scores were calculated with the “CONACC” mode. The phylogenetic tree was constructed through referring published literature and the divergence times among species were obtained from TimeTree [8].

**Comparison of the quality of identified conserved elements in *A. thaliana***

The conserved elements of *A. thaliana* identified in this work were compared with those from Haudry *et al.*’s work [9]. As the conserved elements in coding regions were not released in Haudry *et al.*’s work [9], we only compared the conserved elements in non-coding regions here. The multiple genome alignments and coordinates of conserved noncoding sequences for *A. thaliana* from Haudry *et al.*’s work were downloaded from <http://mustang.biol.mcgill.ca:8885/> [9]. The overlapping analyses were performed between Haudry *et al.*’s work and this work in the aspects of aligned regions and conserved regions, respectively. The genomic regions aligned in both of the two datasets were divided into four classes: ‘Neither’ (nonconserved in both of the two dataset), ‘Both’ (conserved in both of the two datasets), ‘Haudry *et al.*’ (only conserved in Haudry *et al.*’s dataset) and ‘this work’ (only conserved in our dataset). Then, the average ratio of identical bases was defined to assess the quality of identified conserved elements. For each aligned base, the ratio of identical bases means the percentage of the aligned query species that showed the identical base as the target species (*A. thaliana*). Wilcoxon rank sum test was performed to test whether there was significant difference in the ratio of identical bases between those two classes of genomic regions (‘Haudry *et al.*’ and ‘This work’).

**Comparison of conservation landscapes across kingdoms**

For the 63 plant species, the gene annotations and conserved elements mentioned above (Section ‘the workflow for establishing conservation landscapes across green plants’) were used. Corresponding datasets for the other four non-plant species were downloaded from UCSC genome browser [5] (tracks ‘ncbiRefSeq’ and ‘phastConsElements’ respectively). The proportion of non-coding regions in the genome, proportion of coding regions and non-coding regions conserved were calculated based on the gene annotation and conserved elements mentioned here.

**Comparison of transcriptional signals between conserved and nonconserved sequences in unannotated intergenic regions**

The 113 RNA-seq datasets in *A. thaliana* were downloaded from the Araport11 [10]. Reads were mapped to the *Arabidopsis* genome (TAIR10) [11] using Hisat2 (v2.1.0) [12] with the guide of Araport11 gene annotation [10]. Unannotated Intergenic regions were defined as the regions not overlap with gene bodies or their 5kb flanking regions, and they were further divided into conserved unannotated regions and nonconserved unannotated regions based on whether overlapped with conserved elements identified in *A. thaliana*. The read depth for each base pair of the two classes was obtained for each dataset using Bedtools (v2.26.0) [13]. Regions with a read depth no less than five and supported by at least three datasets were assigned as actively transcriptional regions. The actively transcriptional ratio means the proportion of actively transcriptional regions in the entire region. To compare the expression levels in the actively transcriptional regions between these two classes of intergenic regions, the read depths in the regions showing expression signals (with a read depth no less than 5 in at least 3 samples) were calculated based on the mapping results.

**Inference of RNA secondary structures**

The transcript sequences (cDNA) for *A. thaliana* were downloaded from Araport11 [10]. The secondary structure of each transcript was inferred using RNAfold (v2.1.9) [14] with the option of “--noLP”. The positions of stem regions were firstly parsed from the RNAfold prediction result, and then were mapped to the genome coordinates.

**The workflow of FunTFBS to screen for functional TF binding sites and interactions**

For a putative TF binding site (TFBS), the corresponding sequence was firstly extracted from the genome sequence. Then the motif frequency of each base pair in corresponding binding motifs was calculated, and conservation scores (PhyloP) were extracted from conservation landscape established in this work. A Pearson correlation coefficient was calculated between the motif base frequencies and the absolute value of the PhyloP scores. The putative TFBS with significant (p-value <=0.05) and high (Pearson correlation coefficient >=0.5) correlation between motif base frequencies and absolute value of the PhyloP score would be assigned as a functional TFBS (Figure S4). For functional interaction inference, a gene with its promoter region (transcriptional start site (TSS): -500 bp to +100 bp) overlapped with a functional TFBS will be assigned as the target gene of the corresponding TF.

**Generation of the evaluation dataset for functional TF binding site screening using time-course TF binding and expression data**

Song *et al.* generated 114 time-course ChIP-seq datasets (with at least two replicates for each condition; before and after treatment with abscisic acid (ABA)) for 21 TFs and corresponding RNA-seq data in *A. thaliana*, providing an ideal resource to generate a dataset to evaluate the performance of functional TFBS screening methods. For ChIP-seq analysis, the raw data were download from the GEO (GSE80564) [15]. And then the reads were mapped to the TAIR10 genome using bowtie (v1.1.2) [16] with the options of “-m 1 --best --strata -S --chunkmbs 256” and thus only uniquely mapped reads were kept for further analysis. Peaks were called for each TF in each condition using MACS2 [17] with the options of “--nomodel --extsize 250”. The peaks with p-value < 10^-5^ were then classified into two classes, “Less reliable” and “Highly reliable” peaks, based on the consistency among replicates from the IDR pipeline (cutoff: 0.01) [18]. Accordingly, TFBS overlapped with “Less reliable” and “Highly reliable” peaks were assigned as “Less reliable” and “Highly reliable” TFBS, respectively. To identify the functional TFBSs (differential binding will result in differential expression of the target genes), the differential binding peaks (defined as “ABA-response peaks”) between the two conditions (‘normal’ and ‘ABA-treated for 4 hour’) were firstly identified using DiffBind (v2.2.5) [19] with the “Highly reliable” peaks as input. Then, the gene expression data for the corresponding conditions were downloaded from the GEO (<ftp://ftp.ncbi.nlm.nih.gov/geo/series/GSE80nnn/GSE80565/suppl/GSE80565_RNAseq_ABAtimeSeries_rpkm_log2FC_FDR_filterCPM2n2.txt.gz>) [15] and the differentially expressed genes between normal and ‘ABA-treated for 4 hour’ were extracted from the gene expression data (cutoff: FDR <= 0.01). TFBSs in the “ABA-response peaks” with corresponding gene differentially expressed were assigned as “Functional” TFBSs.

**Evaluation of the performance of functional TF binding site screening methods based on the evaluation dataset**

Motifs directly called using the ChIP-seq dataset mentioned above were downloaded from the website (<http://www.abatf.net/>) [15]. For TFs with more than one motif, only the most significant one for each TF was kept for the following analyses. The TFBSs were scanned by the motifs in the gene promoter regions (TSS: -500 bp to +100 bp) using FIMO (v4.10.0) [20] (cutoff: p value <=1e-5) (defined as ‘putative TFBSs’).

To check whether the FunTFBS algorithm could distinguish functional TFBSs from non-functional ones, the putative TFBSs from directly motif scanning were classified into “Consistent” and “Inconsistent” TFBSs based on whether they passed the correlation test of FunTFBS or not. Then, the putative TFBSs were further screened by DNase-seq footprints, conserved elements, and evolutionary footprints (FunTFBS) for functional TFBSs as follows: 1) DNase-seq footprint-based method: the DNase-seq footprint data in *A. thaliana* were downloaded from PlantTFDB [1] and merged all tissues and conditions together. The putative TFBSs with at least 50% overlapped with merged DNase-seq footprints were assigned as “DNase-seq footprint TFBS (DF TFBSs)”; 2) Conserved element-based method: The putative TFBSs with at least 50% overlapped with conserved elements established in this work were assigned as “Conserved TFBSs”; 3) Evolutionary footprint-based method (FunTFBS): the putative TFBSs that passed the screening of FunTFBS method were assigned as “FunTFBSs”.

When comparing “Consistent TFBSs” with “Inconsistent TFBSs” and comparing “FunTFBSs” with ‘DF TFBSs’ and 'Conserved TFBSs’, the percentage of putative TFBSs supported by the three classes of evaluation dataset were used as the corresponding background, respectively. Compared with background, the fold change of the percentage of screened TFBSs supported by evaluation dataset were calculated to assess the performance of different screening methods. 1000 times of subsampling (sampling 50% TFBSs with replacement) were performed to check the significant and robust of the comparison, and the p-values were calculated based on the 1000 subsampling results. For example, when checking whether “Consistent TFBSs” shows higher percentage of putative TFBSs supported by “Functional” TFBSs” compared with “Inconsistent TFBSs”, the p-value was calculated as the proportion of samplings where the hypothesis is rejected.

**Evaluation of the performance of functional TFBS screening methods in inferring functional transcriptional interactions**

The putative TFBSs from directly motif scanning for 599 TFs of *A. thaliana* were downloaded from PlantRegMap [1]. The putative TFBSs were further screened by the three functional TFBS screening methods as described in the section “Evaluation of the performance of functional TF binding site screening methods based on the evaluation dataset”. The resulted TFBSs were used to infer transcriptional regulatory interactions between TFs and target genes based on whether one or more TFBSs of a TF were located in the promoter region of target genes. We evaluated the performance of these methods based on three different features as follows:

1) Evaluation based on published functional transcriptional regulatory interactions. We downloaded the manual-curated, functional-confirmed interactions from *Arabidopsis* transcriptional regulatory map (ATRM) [21]. Only TFs with a binding motif available were kept for evaluation. The percentage of interactions, inferred by directly motif scanning, supported by ATRM was used as the background. Compared with background, the fold changes in the percentage of screened interactions supported by ATRM were calculated to evaluate the three methods. 1000 subsampling (sampling 50% TFBSs with replacement) were performed to check the significant and robust of the comparison, and the p-values were calculated based on the 1000 sampling results.
2) Evaluation based on the percentage of regulatory pairs that coexist in the same biological process [22]. The GO slim data (<https://www.arabidopsis.org/download_files/Public_Data_Releases/TAIR_Data_20140630/ATH_GO_GOSLIM.txt>) for *A. thaliana* was downloaded from TAIR10 [11]. Only the GO “Biological Process” annotations with experimental evidences (evidence code: IDA, IPI, IMP, IGI, or IEP) were used and GO terms commonly shared by TFs (e.g., “GO:0003700: DNA-binding transcription factor activity”) were removed for following analysis. Regulatory pair co-existing in the same biology processes means a TF and its target gene sharing one or more “Biological processes”. The fold changes compared with background, inferred by the putative TFBSs from directly motif scanning, were calculated, and Fisher’s exact tests were performed to assess the interactions inferred by FunTFBSs with the other two methods.
3) Evaluation based on the percentage of regulatory pairs that are highly correlated in expression. The Pearson correlation coefficient data among *A. thaliana* genes in expression was downloaded from ATTED-II [23]. Gene pairs with Pearson correlation coefficient no less than 0.3 were assigned as highly correlated in expression here. The percentage of regulatory pairs inferred by the putative TFBSs from directly motif scanning were used as background, and the fold changes compared with background were calculated. Fisher’s exact tests were performed to assess the interactions inferred by FunTFBSs with the other two methods.

**Identification of putative expression quantitative trait loci (eQTLs) using the data from 1001 Genome Project**

The genomic variation data of 1,135 individuals in *A. thaliana* and corresponding accession information were downloaded from 1001 Genomes Project (genomic variation: <http://1001genomes.org/data/GMI-MPI/releases/v3.1/1001genomes_snp-short-indel_only_ACGTN.vcf.gz>; accession information: <https://1001genomes.org/accessions.html>) [24]. The gene expression matrix for 727 individuals was downloaded from the GEO (<ftp://ftp.ncbi.nlm.nih.gov/geo/series/GSE80nnn/GSE80744/suppl/GSE80744_ath1001_tx_norm_2016-04-21-UQ_gNorm_normCounts_k4.tsv.gz>) [25]. Only the 665 individuals with both the variation data and expression data were kept for further analysis. Then, Matrix eQTL [26] was employed to identify *cis-*eQTLs with country, latitude and longitude as the covariates (cutoff: BH-corrected p value < 0.01). Fisher’s exact test was performed to test whether eQTLs were enriched in FunTFBSs compared with non-FunTFBS regions in the promoters of genes, and Wilcoxon rank sum test was performed to test whether there was significant difference in the expression of the corresponding genes presenting the reference allele and the alternative allele.

**GO enrichment analysis for the targets genes inferred by FunTFBS**

GO enrichment analysis for the target genes inferred by FunTFBS was performed using the GO enrichment tool in PlantRegMap [1]. Default background was used and only the terms with p-values (BH-corrected) less than 0.05 were shown.

**Supplementary Figures**

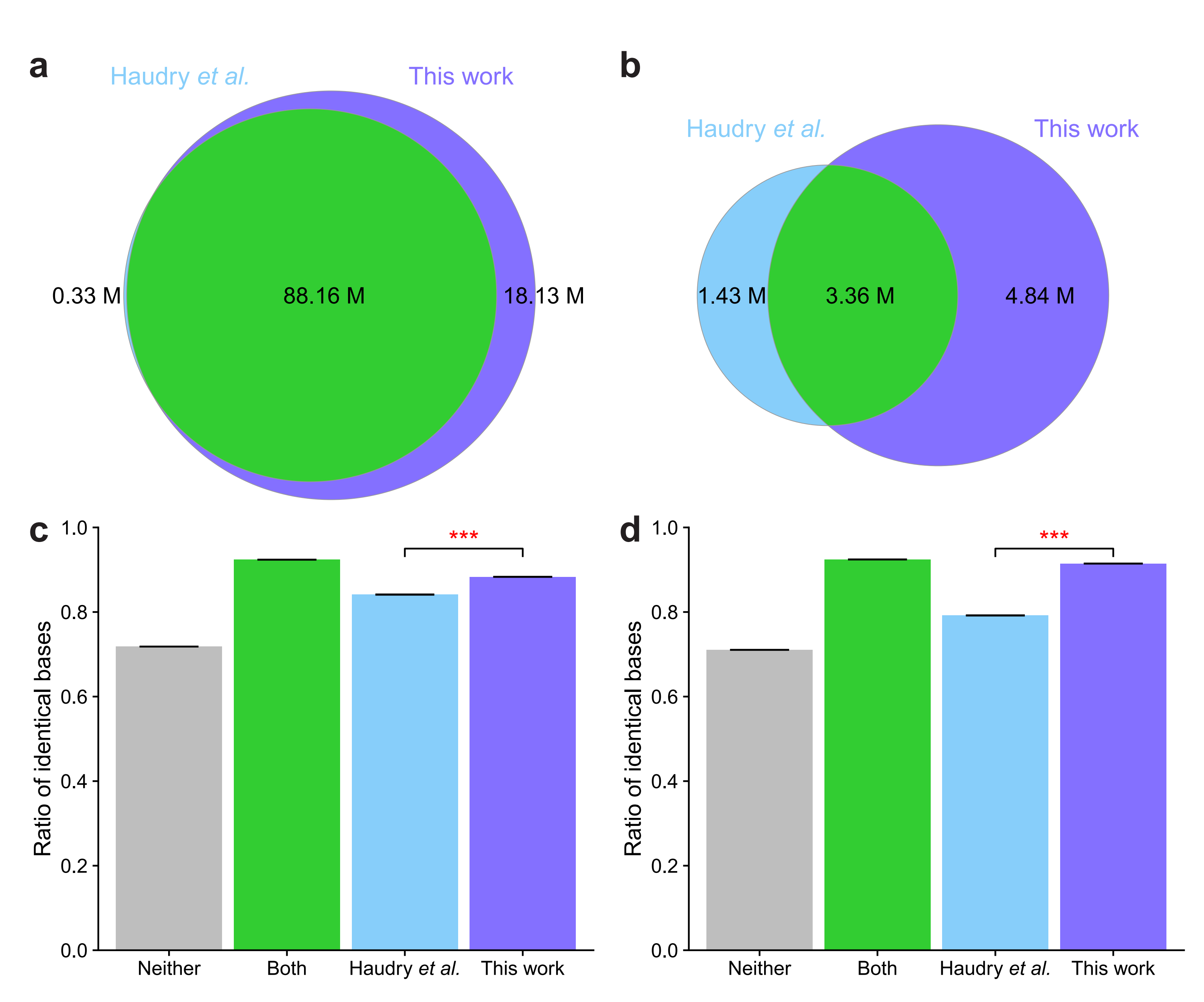

**Figure S1.** Comparison of conserved non-coding sequences in *A. thaliana* identified by this work with those from Haudry *et al.*’s study [9]. **a,b,** Overlapping of genome sequences aligned (**a**) and conserved in non-coding regions (**b**) between this work and that of Haudry *et al.*’s. **c,d,** Quality evaluation of identified conserved non-coding sequences using ratio of identical bases in aligned species for all bases in the conserved non-coding regions, based on the multiple genome alignments from Haudry *et al.*’s study (**c**) and that of this work (**d**). The higher in the ratio of identical bases indicates a higher quality of identified conserved sequences. (Error bar indicates 95% confidence interval; Wilcoxon rank sum test, *** p-value < 0.001.)

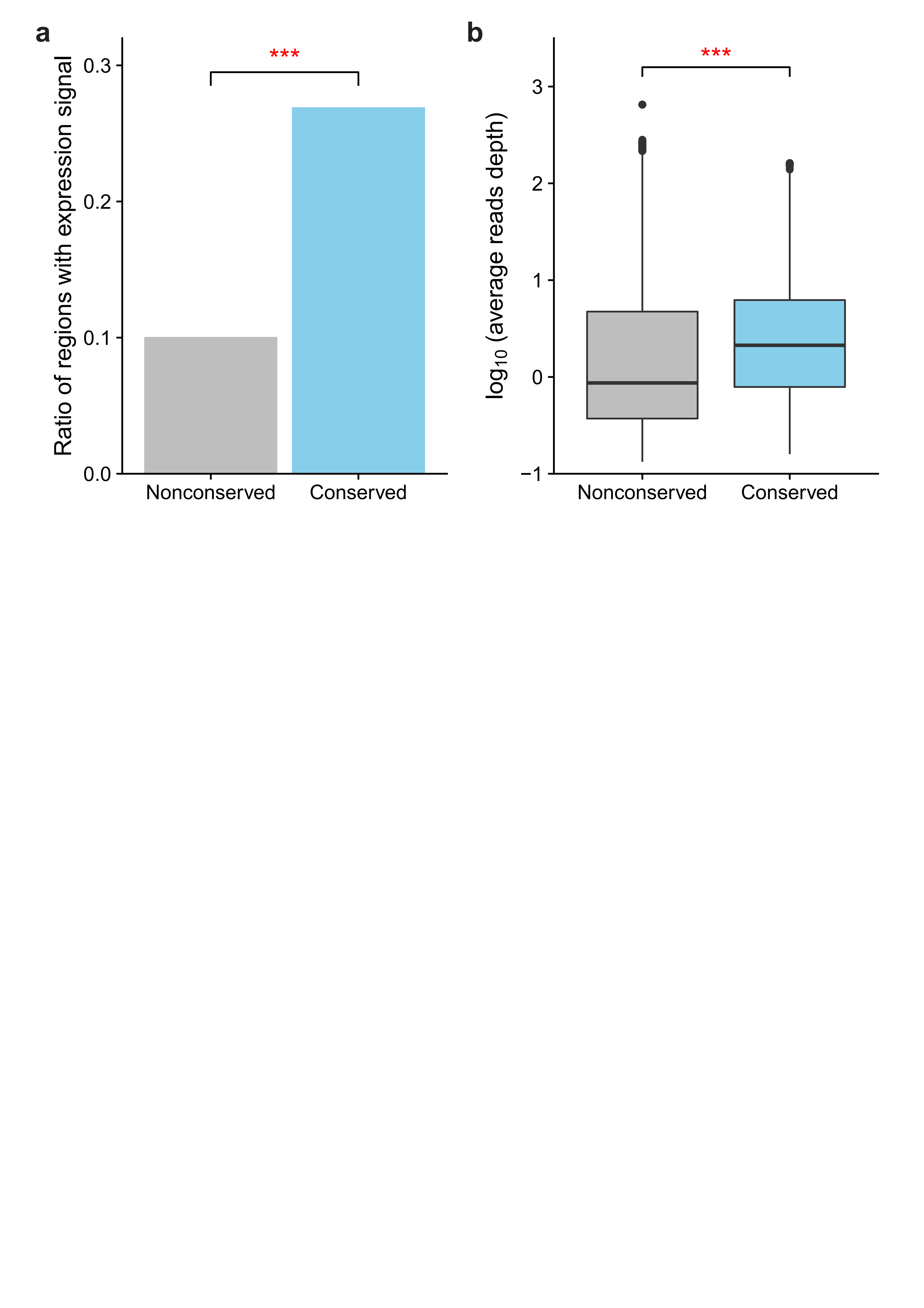

**Figure S2.** Comparison of the expression signal between nonconserved and conserved sequences in the intergenic regions of *A. thaliana*. **a,** The ratio of regions with expression signal in the nonconserved and conserved sequences (Fisher’s exact test, *** p-value < 0.001). **b,** The average expression level of regions with expression signal in the nonconserved and conserved sequences. (Wilcoxon rank sum test, *** p-value < 0.001.)

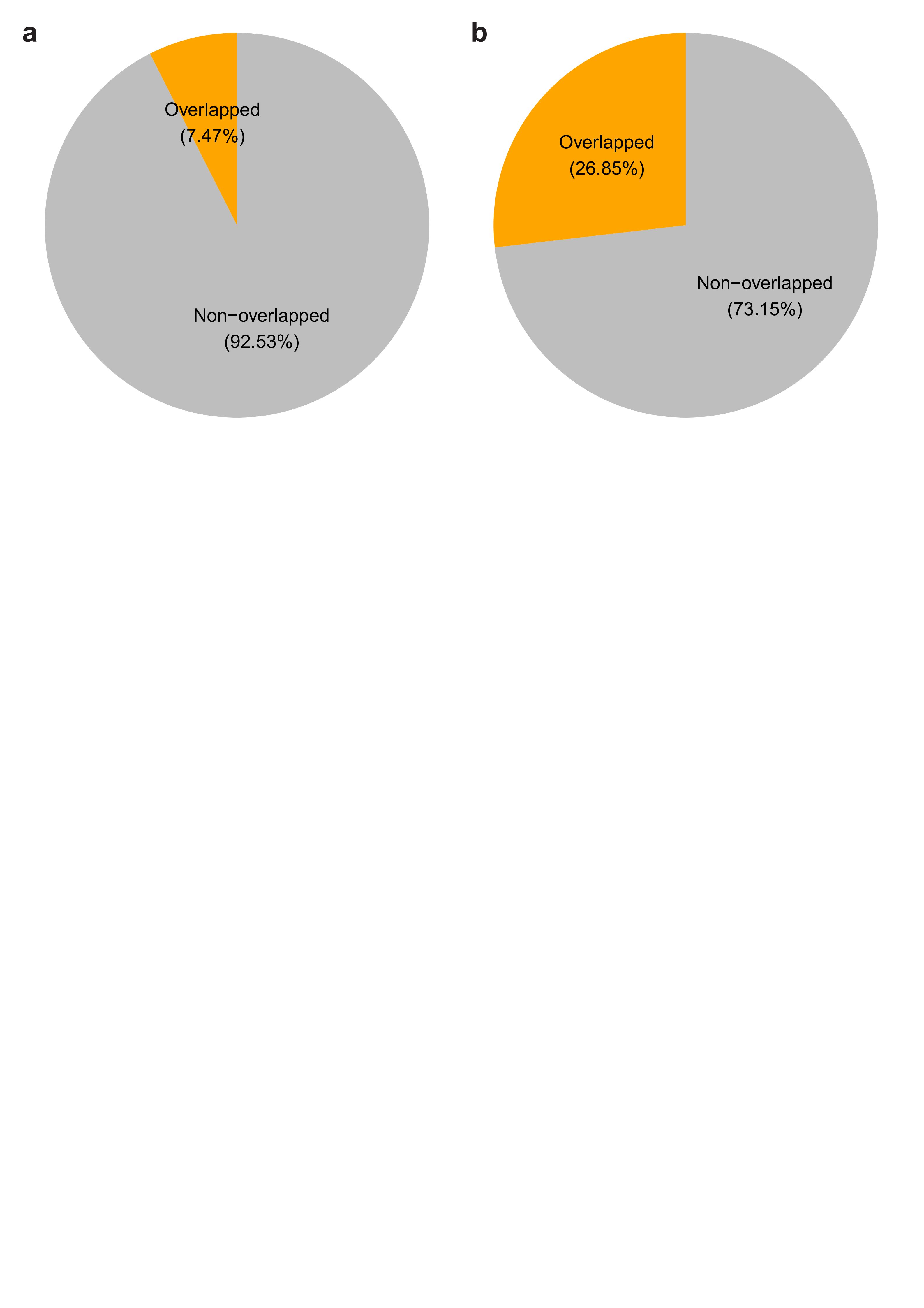

**Figure S3.** Other functional elements overlapped with conserved elements in the gene promoter regions of *A. thaliana*. **a,** Proportion of noncoding conserved elements in the gene promoter regions that are overlapped with other noncoding RNAs (ncRNAs). **b,** Proportion of noncoding conserved elements in the gene promoter regions that are overlapped with the stem regions of predicted RNA secondary structures.

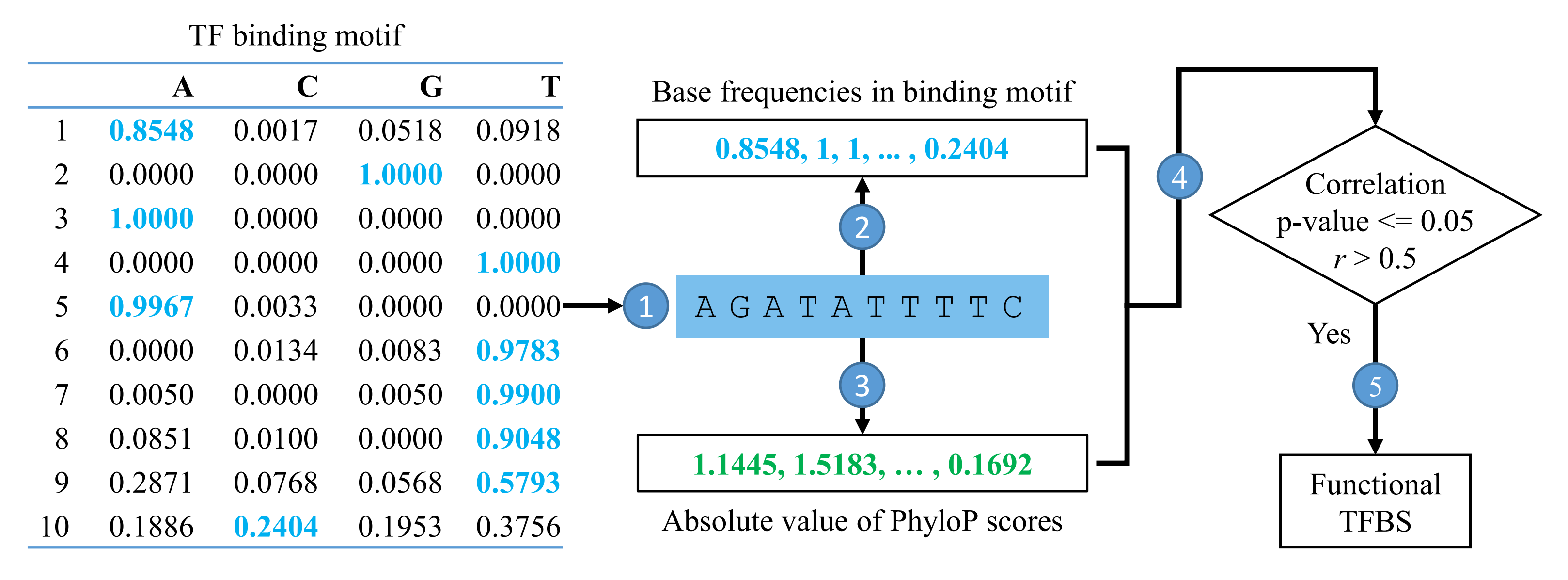

**Figure S4.** A case showing the workflow of the FunTFBS tool to screen for functional transcription factor binding sites (TFBSs). For a TFBS candidate, the genomic sequences will be extracted **(1)**, and the base frequency in the binding motifs **(2)** and the PhyloP score **(3)** for each base of TFBS will be calculated. Then, the Pearson correlation coefficient between base frequencies in binding motifs and absolute value of PhyloP scores will be calculated **(4)** and only the TFBS candidate with significant correlation (p-value <= 0.05) and correlation coefficient greater than 0.5 is kept as a functional TFBS **(5)**.

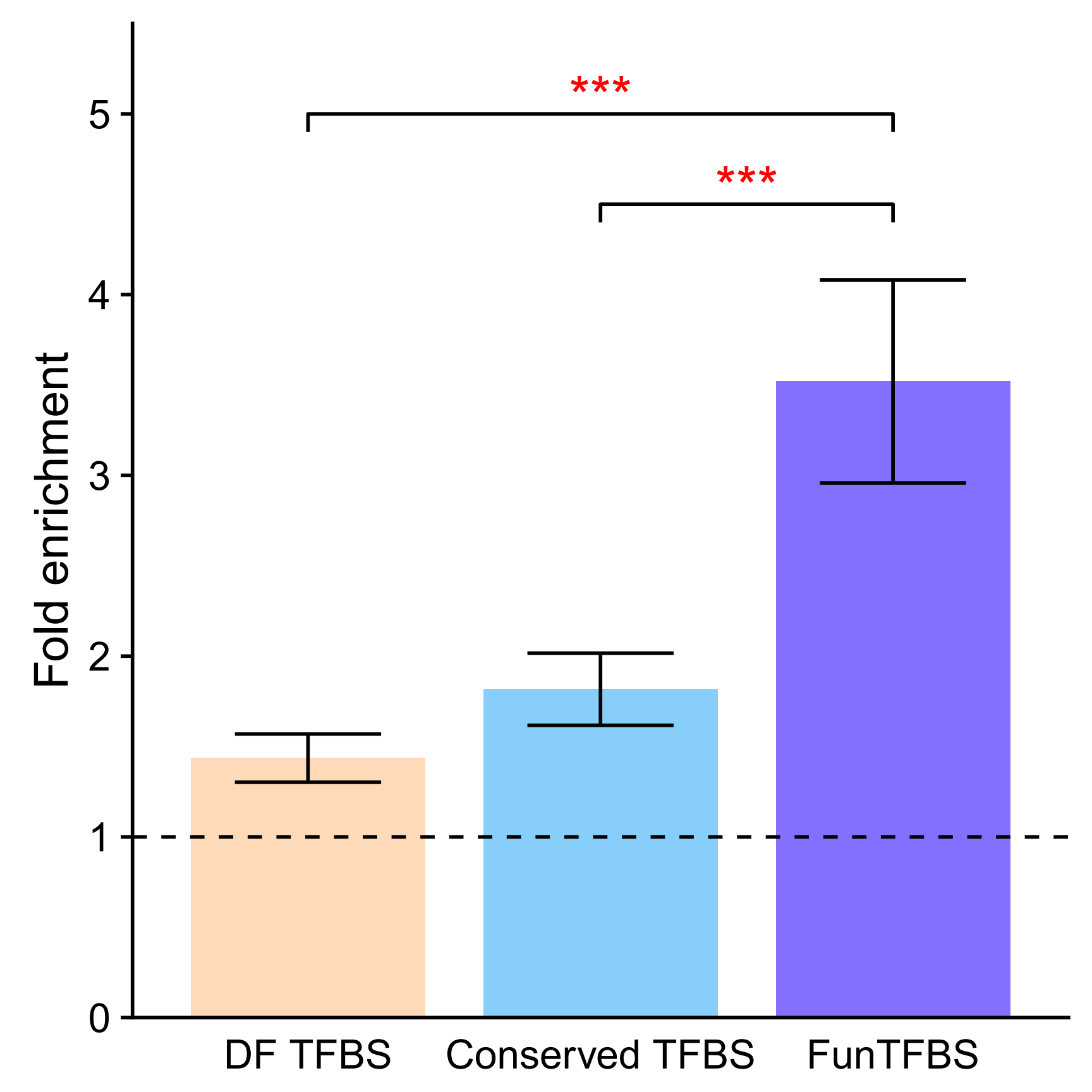

**Figure S5.** Evaluation of the regulations inferred by DNase-seq footprint (DF) TFBS, Conserved TFBS and FunTFBS based on the functional regulations from *Arabidopsis* transcriptional regulatory map (ATRM). Fold changes of the percentage of regulations that were supported by the ATRM data are shown, as inferred by the kinds of methods compared with the regulations directly from motif scanning. (Error bar indicates standard deviation of the average fold changes from 1000 subsampling; p-value was calculated based on the results of the 1000 subsampling, *** p-value < 0.001.)

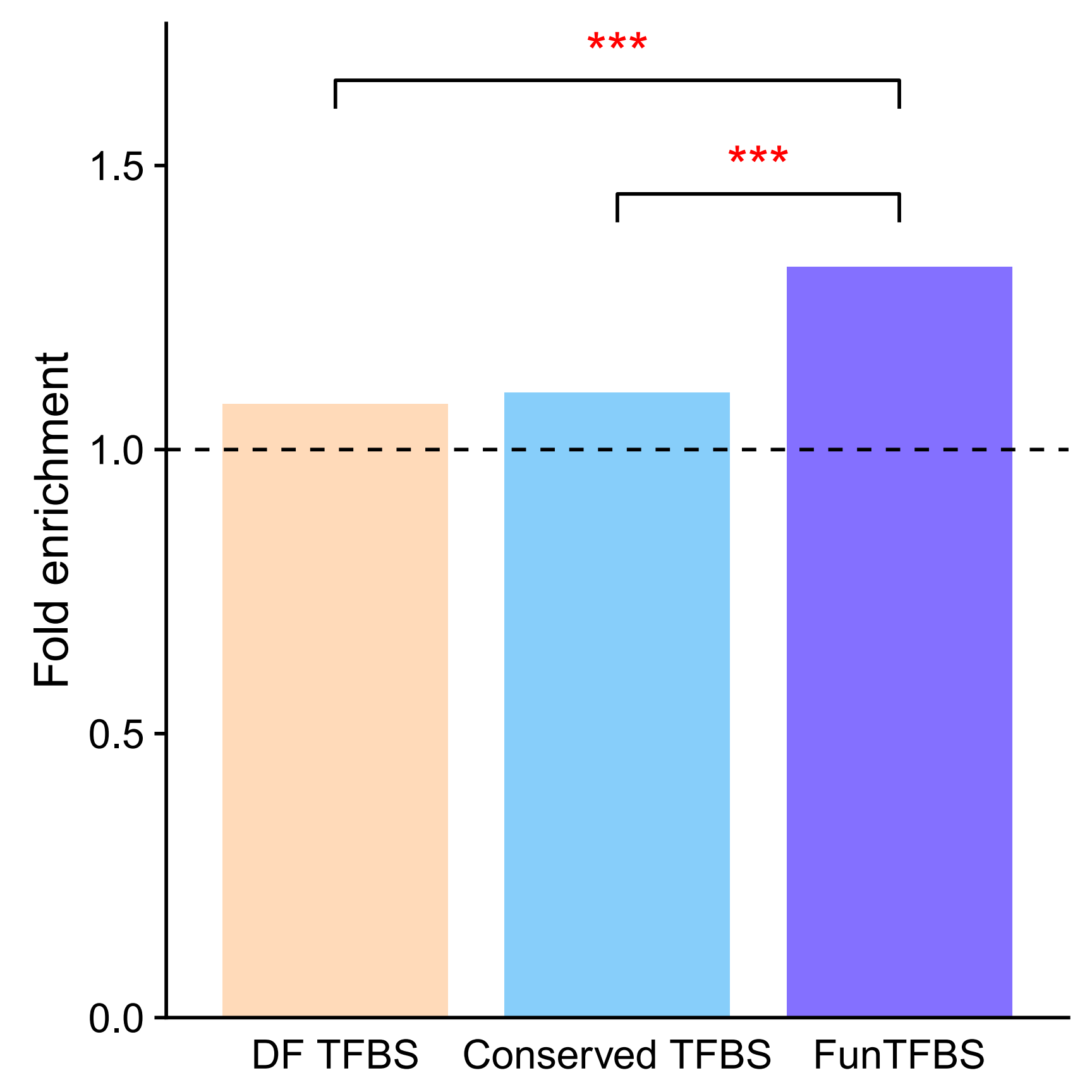

**Figure S6.** Evaluation of the regulations inferred by DNase-seq footprint (DF) TFBS, Conserved TFBS and FunTFBS based on the percentage of regulatory pairs that coexist in the same biological process. Fold changes of percentage of regulations coexisting in the same GO biological process are shown, as inferred by the kinds of methods compared with the regulation directly from motif scanning. (Fisher’s exact test, *** p-value < 0.001.)

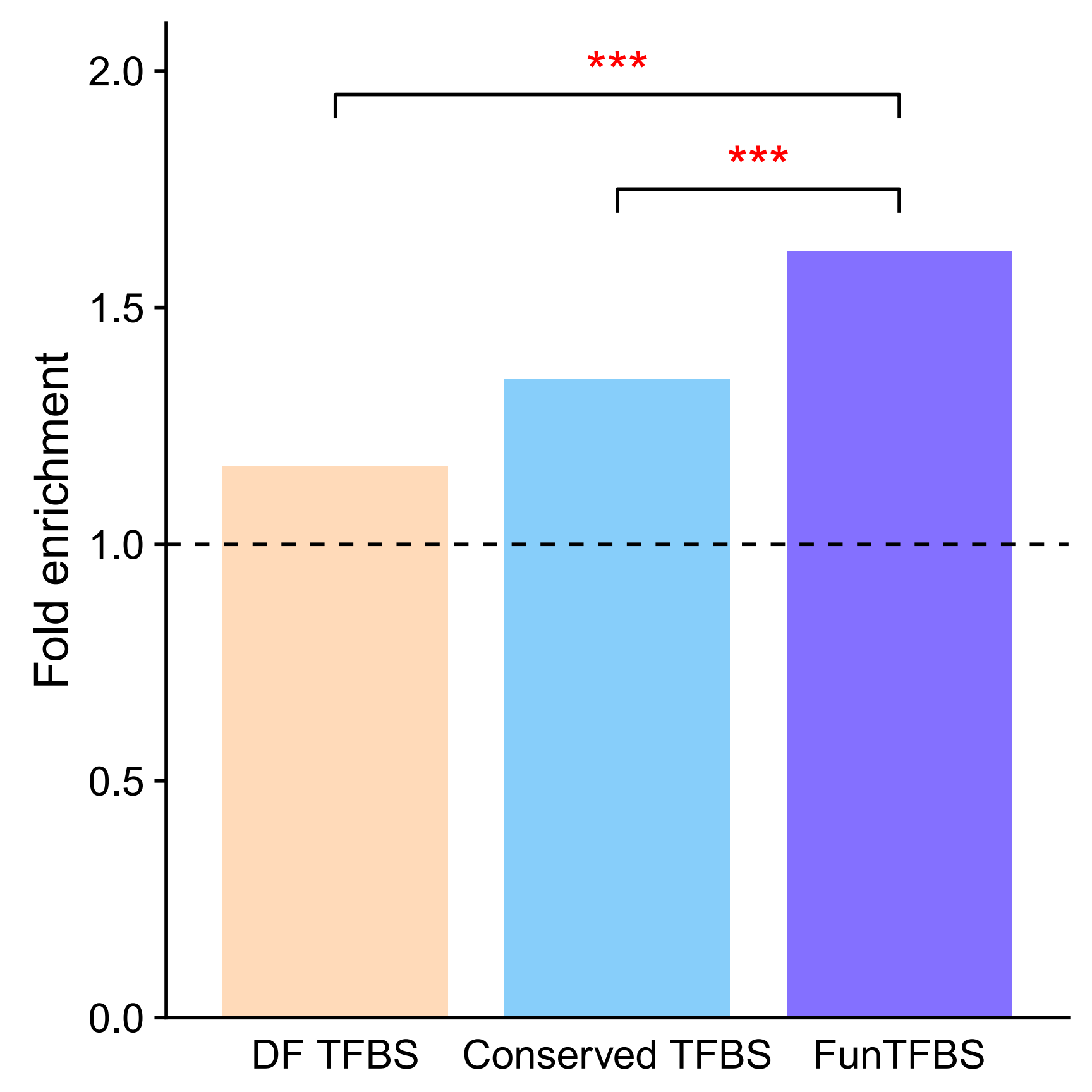

**Figure S7.** Evaluation of the regulations inferred by DNase-seq footprint (DF) TFBS, Conserved TFBS and FunTFBS based on the percentage of regulatory pairs that are highly correlated in expression. Fold changes of the percentage of regulations with a high expression correlation (Pearson correlation coefficient *r* >= 0.3) are shown, as inferred by the kinds of methods compared with the regulation directly from motif scanning. (Fisher’s exact test, *** p-value < 0.001.)

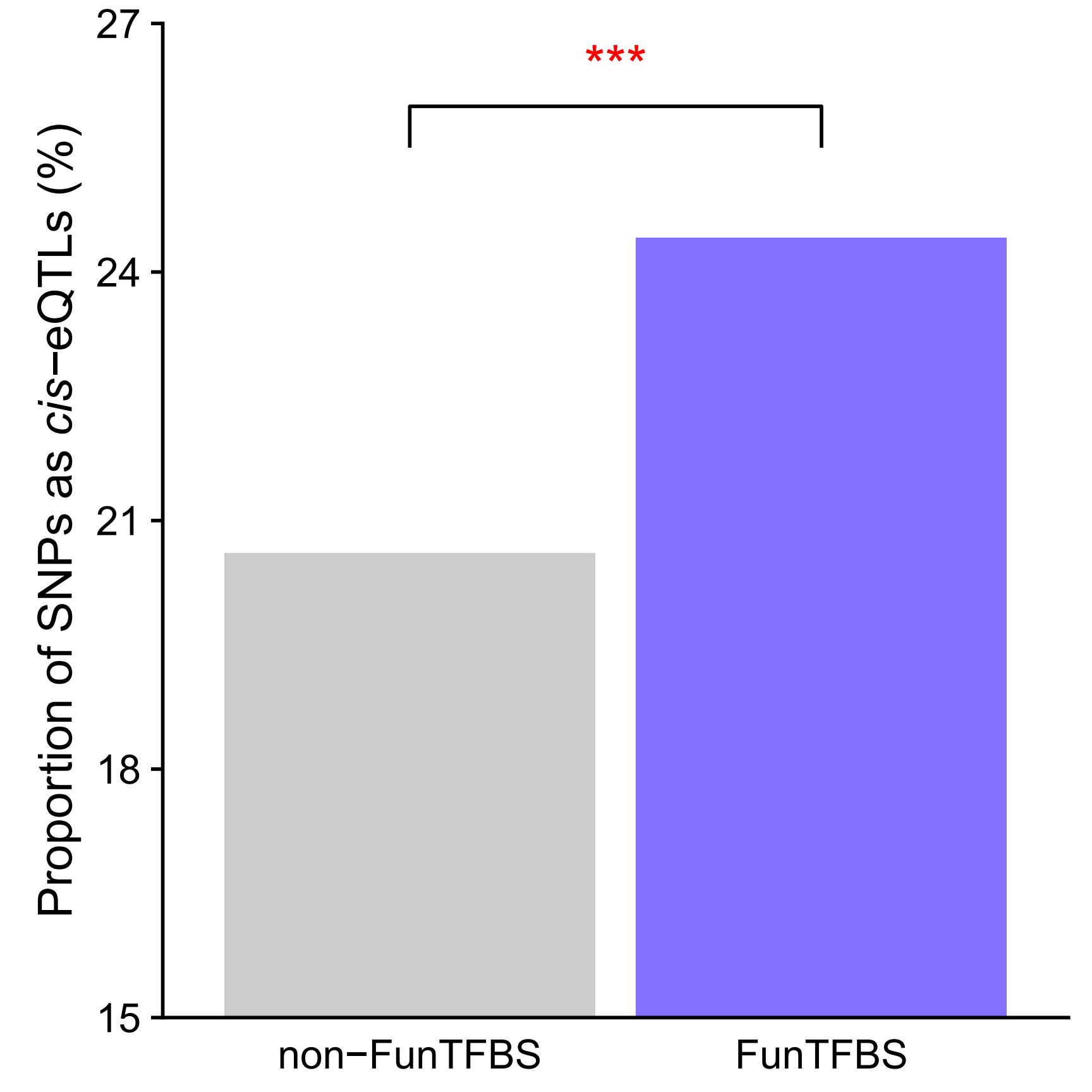

**Figure S8.** The proportion of SNPs identified as *cis-*expression quantitative trait loci (*cis-*eQTLs) for the SNPs located in non-FunTFBS (Inconsistent TFBS) and FunTFBS (Consistent TFBS) regions (Fisher’s exact test, *** p-value < 0.001.)

**Supplementary Tables**

**Table S1.** Summary of data sources and results of multiple genome alignments

| Group | Species | Genome assembly | Genome size (Mb) | Aligned ratio in CDS (%) | Conserved ratio in aligned CDS (%) | Aligned ratio in genome (%) | Conserved ratio in aligned genome (%) |
| --- | --- | --- | --- | --- | --- | --- | --- |
| BOP clade | *Brachypodium distachyon* | JGI(v3.1) | 271.16 | 97.29 | 54.40 | 67.77 | 22.64 |
|  | *Brachypodium stacei* | JGI(v1.1) | 233.92 | 98.53 | 56.30 | 81.28 | 21.19 |
|  | *Hordeum vulgare* | IBSC(v1.0) | 4,014.49 | 98.86 | 65.50 | 16.24 | 7.60 |
|  | *Leersia perrieri* | OGE(v1.4) | 266.69 | 99.61 | 58.40 | 95.26 | 15.87 |
|  | *Oryza brachyantha* | OGE(v1.4b) | 258.58 | 97.91 | 63.50 | 89.61 | 16.58 |
|  | *Oryza glaberrima* | AGI(v1.1) | 310.25 | 99.59 | 59.60 | 95.59 | 15.99 |
|  | *Oryza longistaminata* | BGI(v1.0) | 291.29 | 99.85 | 56.10 | 88.87 | 14.97 |
|  | *Oryza sativa subsp. japonica* | MSU(v7) | 374.47 | 99.05 | 56.30 | 86.06 | 17.65 |
|  | *Phyllostachys heterocycla* | ICBR(v1.0) | 1,750.71 | 98.37 | 67.20 | 73.87 | 7.76 |
| Brassicales | *Aethionema arabicum* | VEGI(v2.5) | 191.05 | 99.62 | 58.30 | 81.88 | 17.72 |
|  | *Arabidopsis halleri* | JGI(v1.1) | 104.56 | 99.32 | 58.50 | 78.74 | 27.16 |
|  | *Arabidopsis lyrata* | JGI(v1.0) | 204.15 | 99.36 | 54.40 | 83.60 | 17.90 |
|  | *Arabidopsis thaliana* | TAIR10 | 119.67 | 99.68 | 65.20 | 89.23 | 29.16 |
|  | *Arabis alpina* | MPIPBR(v4) | 279.69 | 98.10 | 60.70 | 87.33 | 15.27 |
|  | *Boechera stricta* | JGI(v1.2) | 185.28 | 99.76 | 62.00 | 84.79 | 19.84 |
|  | *Brassica napus* | Genoscope(v4.1) | 850.29 | 99.91 | 68.30 | 86.40 | 17.30 |
|  | *Brassica oleracea* | BOL | 470.37 | 99.86 | 52.70 | 90.54 | 14.95 |
|  | *Brassica rapa* | Brapa_1.0 | 274.16 | 99.85 | 53.40 | 95.17 | 18.62 |
|  | *Camelina sativa* | CSGP(v2) | 622.93 | 99.72 | 54.90 | 91.66 | 16.52 |
|  | *Capsella grandiflora* | JGI(v1.1) | 96.81 | 99.93 | 54.30 | 88.58 | 27.18 |
|  | *Capsella rubella* | JGI(v1.0) | 132.26 | 99.89 | 54.00 | 96.47 | 20.81 |
|  | *Carica papaya* | ASGPB(v0.4) | 327.73 | 87.25 | 64.40 | 30.68 | 28.71 |
|  | *Eutrema salsugineum* | JGI(v1.0) | 240.54 | 99.76 | 68.10 | 94.70 | 15.83 |
|  | *Raphanus sativus* | RGD(v1.0) | 314.68 | 99.70 | 55.20 | 74.79 | 19.82 |
|  | *Sisymbrium irio* | VEGI(v1) | 229.50 | 98.24 | 53.50 | 79.04 | 24.26 |
|  | *Tarenaya hassleriana* | ASM46358v1 | 242.74 | 99.01 | 64.20 | 76.21 | 21.32 |
|  | *Thellungiella parvula* | thellungiella(v2.0) | 118.66 | 98.90 | 67.30 | 96.26 | 26.74 |
| Fabales | *Arachis duranensis* | NCGR_PGC(v1) | 1,084.24 | 99.70 | 60.40 | 85.81 | 7.01 |
|  | *Arachis ipaensis* | NCGR_PGC(v1) | 1,353.83 | 99.64 | 64.50 | 91.91 | 12.65 |
|  | *Cajanus cajan* | IIPG(v1) | 564.39 | 99.66 | 54.10 | 89.08 | 8.64 |
|  | *Cicer arietinum* | ASM33114v1 | 518.13 | 99.77 | 65.70 | 82.06 | 8.80 |
|  | *Glycine max* | Wm82.a2.v1 | 974.36 | 99.51 | 54.90 | 88.23 | 14.95 |
|  | *Glycine soja* | TCUHK(v1.0) | 839.51 | 99.93 | 55.20 | 93.13 | 10.39 |
|  | *Lotus japonicus* | Kazusa(v3.0) | 447.42 | 96.57 | 60.40 | 79.73 | 19.34 |
|  | *Medicago truncatula* | Mt4.0v1 | 407.88 | 95.24 | 56.20 | 77.61 | 14.15 |
|  | *Phaseolus vulgaris* | JGI(v1.0) | 519.13 | 99.81 | 60.60 | 85.41 | 8.49 |
|  | *Trifolium pratense* | JGI(v2) | 292.06 | 97.95 | 61.70 | 85.34 | 15.06 |
|  | *Vigna angularis* | SNU(v3) | 438.08 | 99.69 | 53.40 | 88.16 | 9.21 |
|  | *Vigna radiata* | SNU(v6) | 458.43 | 99.85 | 54.00 | 91.53 | 8.18 |
| Lamiales | *Dorcoceras hygrometricum* | CNU(v1) | 1,193.68 | 87.38 | 62.50 | 46.97 | 21.92 |
|  | *Mimulus guttatus* | JGI(v2.0) | 306.80 | 96.38 | 65.90 | 58.38 | 20.85 |
|  | *Sesamum indicum* | S_indicum_v1.0 | 266.94 | 94.26 | 66.90 | 53.08 | 22.27 |
|  | *Utricularia gibba* | UGSP(v4.1) | 74.04 | 84.90 | 59.70 | 57.35 | 39.96 |
| Malpighiales | *Linum usitatissimum* | BGI(v1.0) | 295.20 | 92.29 | 63.30 | 55.07 | 29.24 |
|  | *Manihot esculenta* | JGI(v6.1) | 576.56 | 97.31 | 63.90 | 58.95 | 13.62 |
|  | *Populus trichocarpa* | JGI(v3.0) | 429.86 | 97.90 | 65.20 | 71.79 | 14.10 |
|  | *Ricinus communis* | JCVI(v0.1) | 317.70 | 96.68 | 64.30 | 70.27 | 13.76 |
|  | *Salix purpurea* | JGI(v1.0) | 457.63 | 99.80 | 60.00 | 69.02 | 13.82 |
| PACMAD clade | *Dichanthelium oligosanthes* | Dichan(v1) | 566.99 | 99.69 | 60.70 | 73.91 | 9.56 |
|  | *Eragrostis tef* | Tef(v1.1.2) | 588.97 | 98.99 | 58.70 | 71.31 | 13.39 |
|  | *Oropetium thomaeum* | JGI(v1.0) | 243.17 | 99.51 | 62.10 | 94.18 | 12.29 |
|  | *Panicum virgatum* | JGI(v3.1) | 1,625.16 | 98.75 | 53.70 | 64.33 | 11.67 |
|  | *Setaria italica* | JGI(v2.2) | 404.50 | 99.51 | 57.00 | 93.25 | 13.57 |
|  | *Setaria viridis* | JGI(v1.1) | 392.92 | 99.44 | 59.00 | 88.15 | 15.37 |
|  | *Sorghum bicolor* | JGI(v3.1) | 729.66 | 98.07 | 61.30 | 32.44 | 27.75 |
|  | *Zea mays* | AGPv3 | 2,067.23 | 95.14 | 61.50 | 20.54 | 13.12 |
|  | *Zoysia pacifica* | ZGD(v1.0) | 382.11 | 96.97 | 55.30 | 90.83 | 18.52 |
| Rosales | *Fragaria vesca* | GDR(v1.1) | 206.89 | 90.45 | 56.60 | 50.37 | 24.83 |
|  | *Morus notabilis* | BGI(ASM41409v2) | 298.48 | 92.80 | 66.60 | 58.12 | 19.88 |
|  | *Prunus mume* | BGI(v1.0) | 228.49 | 99.92 | 56.20 | 91.45 | 19.91 |
|  | *Prunus persica* | JGI(v2.1) | 226.73 | 99.35 | 54.40 | 92.67 | 14.16 |
|  | *Pyrus bretschneideri* | CPETR(v1.0) | 510.29 | 96.29 | 56.50 | 62.92 | 17.47 |
|  | *Ziziphus jujuba* | ZizJuj_1.1 | 425.78 | 98.10 | 63.10 | 66.86 | 18.42 |

**Table S2.** Summary of evolutionary footprints of 63 plant species

| Group | Species | Region with Conservation scores (Mb) | Number of conserved elements | Length of conserved elements (Mb) | Length of conserved CDS (Mb) | Length of conserved non-CDS (Mb) | Conserved in genome (%) | Conserved in CDS (%) | Conserved in non-CDS (%) |
| --- | --- | --- | --- | --- | --- | --- | --- | --- | --- |
| BOP clade | *Brachypodium distachyon* | 183.76 | 421,923 | 41.62 | 20.66 | 20.96 | 15.35 | 52.97 | 9.03 |
|  | *Brachypodium stacei* | 190.12 | 439,851 | 40.30 | 20.05 | 20.25 | 17.23 | 55.52 | 10.24 |
|  | *Hordeum vulgare* | 652.09 | 802,036 | 49.60 | 17.71 | 31.89 | 1.24 | 64.84 | 0.80 |
|  | *Leersia perrieri* | 254.05 | 590,899 | 40.34 | 21.45 | 18.89 | 15.13 | 58.25 | 8.22 |
|  | *Oryza brachyantha* | 231.73 | 543,961 | 38.43 | 20.76 | 17.68 | 14.86 | 62.26 | 7.85 |
|  | *Oryza glaberrima* | 296.59 | 654,090 | 47.45 | 21.22 | 26.22 | 15.29 | 59.38 | 9.55 |
|  | *Oryza longistaminata* | 258.88 | 312,489 | 38.78 | 18.00 | 20.78 | 13.31 | 56.12 | 8.02 |
|  | *Oryza sativa subsp. japonica* | 322.28 | 372,065 | 56.91 | 23.30 | 33.62 | 15.20 | 55.86 | 10.10 |
|  | *Phyllostachys heterocycla* | 1,293.39 | 1,876,004 | 100.44 | 25.26 | 75.18 | 5.74 | 66.15 | 4.39 |
| Brassicales | *Aethionema arabicum* | 156.43 | 1,362,027 | 27.73 | 17.30 | 10.43 | 14.52 | 58.09 | 6.47 |
|  | *Arabidopsis halleri* | 82.34 | 941,813 | 22.37 | 14.60 | 7.76 | 21.39 | 58.17 | 9.77 |
|  | *Arabidopsis lyrata* | 170.68 | 1,242,841 | 30.56 | 19.01 | 11.56 | 14.97 | 54.12 | 6.84 |
|  | *Arabidopsis thaliana* | 106.78 | 1,275,925 | 31.15 | 21.77 | 9.37 | 26.03 | 65.06 | 10.87 |
|  | *Arabis alpina* | 244.26 | 1,863,293 | 37.31 | 14.56 | 22.74 | 13.34 | 59.61 | 8.91 |
|  | *Boechera stricta* | 157.11 | 1,470,668 | 31.17 | 19.69 | 11.49 | 16.82 | 61.95 | 7.48 |
|  | *Brassica napus* | 734.66 | 5,596,980 | 127.15 | 69.11 | 58.05 | 14.95 | 68.31 | 7.75 |
|  | *Brassica oleracea* | 425.89 | 1,280,964 | 63.68 | 27.79 | 35.89 | 13.54 | 52.68 | 8.59 |
|  | *Brassica rapa* | 260.94 | 1,207,474 | 48.60 | 26.53 | 22.07 | 17.73 | 53.35 | 9.83 |
|  | *Camelina sativa* | 571.03 | 4,771,238 | 94.35 | 58.22 | 36.13 | 15.15 | 54.84 | 6.99 |
|  | *Capsella grandiflora* | 85.76 | 1,069,709 | 23.32 | 15.85 | 7.47 | 24.09 | 54.29 | 11.05 |
|  | *Capsella rubella* | 127.60 | 1,047,332 | 26.56 | 17.83 | 8.73 | 20.08 | 53.95 | 8.80 |
|  | *Carica papaya* | 100.55 | 1,477,102 | 28.88 | 13.26 | 15.61 | 8.81 | 56.21 | 5.13 |
|  | *Eutrema salsugineum* | 227.81 | 1,733,740 | 36.07 | 21.96 | 14.11 | 15.00 | 68.03 | 6.77 |
|  | *Raphanus sativus* | 235.37 | 1,103,514 | 46.66 | 23.92 | 22.74 | 14.83 | 55.09 | 8.38 |
|  | *Sisymbrium irio* | 181.42 | 840,003 | 44.03 | 27.75 | 16.28 | 19.18 | 52.65 | 9.21 |
|  | *Tarenaya hassleriana* | 185.00 | 2,725,961 | 39.44 | 22.44 | 17.00 | 16.25 | 63.58 | 8.19 |
|  | *Thellungiella parvula* | 114.23 | 1,224,492 | 30.55 | 20.27 | 10.29 | 25.75 | 66.60 | 11.66 |
| Fabales | *Arachis duranensis* | 930.45 | 989,135 | 65.43 | 24.32 | 40.98 | 6.02 | 60.26 | 3.93 |
|  | *Arachis ipaensis* | 1,244.39 | 1,251,876 | 166.14 | 28.17 | 129.32 | 11.63 | 64.31 | 9.87 |
|  | *Cajanus cajan* | 502.80 | 530,779 | 43.45 | 20.24 | 23.20 | 7.70 | 54.00 | 4.40 |
|  | *Cicer arietinum* | 425.19 | 994,024 | 37.43 | 21.50 | 15.93 | 7.22 | 65.60 | 3.28 |
|  | *Glycine max* | 859.68 | 918,055 | 128.61 | 35.95 | 92.66 | 13.20 | 54.70 | 10.20 |
|  | *Glycine soja* | 781.84 | 1,393,817 | 81.31 | 30.55 | 50.76 | 9.69 | 55.22 | 6.47 |
|  | *Lotus japonicus* | 356.74 | 935,094 | 77.74 | 22.31 | 46.69 | 15.42 | 58.40 | 11.41 |
|  | *Medicago truncatula* | 316.59 | 701,146 | 44.82 | 26.63 | 18.18 | 10.99 | 53.54 | 5.08 |
|  | *Phaseolus vulgaris* | 443.44 | 661,708 | 37.66 | 20.97 | 16.69 | 7.25 | 60.53 | 3.44 |
|  | *Trifolium pratense* | 249.27 | 665,481 | 37.55 | 22.74 | 14.81 | 12.86 | 60.53 | 5.82 |
|  | *Vigna angularis* | 386.23 | 392,458 | 35.58 | 16.65 | 18.92 | 8.12 | 53.27 | 4.65 |
|  | *Vigna radiata* | 419.61 | 544,598 | 34.33 | 15.24 | 19.08 | 7.49 | 53.97 | 4.44 |
| Lamiales | *Dorcoceras hygrometricum* | 560.69 | 5,017,699 | 122.90 | 12.28 | 110.63 | 10.30 | 54.66 | 9.45 |
|  | *Mimulus guttatus* | 179.11 | 543,648 | 37.36 | 21.18 | 16.18 | 12.18 | 63.59 | 5.92 |
|  | *Sesamum indicum* | 141.72 | 364,784 | 31.57 | 20.28 | 11.29 | 11.83 | 63.08 | 4.81 |
|  | *Utricularia gibba* | 42.47 | 280,295 | 16.97 | 13.39 | 3.58 | 22.92 | 50.69 | 7.53 |
| Malpighiales | *Linum usitatissimum* | 162.57 | 1,403,012 | 47.55 | 30.42 | 17.13 | 16.11 | 58.52 | 7.05 |
|  | *Manihot esculenta* | 339.94 | 932,036 | 46.32 | 24.01 | 22.30 | 8.03 | 62.18 | 4.15 |
|  | *Populus trichocarpa* | 308.62 | 572,567 | 43.53 | 30.89 | 12.64 | 10.13 | 63.91 | 3.31 |
|  | *Ricinus communis* | 223.27 | 558,778 | 30.74 | 17.22 | 13.52 | 9.68 | 62.17 | 4.66 |
|  | *Salix purpurea* | 315.86 | 1,093,007 | 43.66 | 27.27 | 16.40 | 9.54 | 59.95 | 3.98 |
| PACMAD clade | *Dichanthelium oligosanthes* | 419.07 | 438,469 | 40.09 | 20.57 | 19.53 | 7.07 | 60.53 | 3.66 |
|  | *Eragrostis tef* | 420.01 | 729,572 | 56.24 | 28.24 | 28.00 | 9.55 | 58.19 | 5.18 |
|  | *Oropetium thomaeum* | 229.04 | 425,813 | 28.15 | 16.64 | 11.51 | 11.58 | 61.88 | 5.32 |
|  | *Panicum virgatum* | 1,045.55 | 1,423,354 | 122.04 | 51.32 | 70.72 | 7.51 | 53.09 | 4.63 |
|  | *Setaria italica* | 377.20 | 715,239 | 51.19 | 23.39 | 27.80 | 12.66 | 56.78 | 7.65 |
|  | *Setaria viridis* | 346.40 | 565,777 | 53.25 | 24.16 | 29.09 | 13.55 | 58.75 | 8.27 |
|  | *Sorghum bicolor* | 236.72 | 561,775 | 65.69 | 23.97 | 41.72 | 9.00 | 60.14 | 6.05 |
|  | *Zea mays* | 424.70 | 772,783 | 55.73 | 25.75 | 29.98 | 2.70 | 58.51 | 1.48 |
|  | *Zoysia pacifica* | 347.07 | 761,736 | 64.30 | 24.03 | 40.27 | 16.83 | 53.64 | 11.94 |
| Rosales | *Fragaria vesca* | 104.22 | 348,338 | 25.88 | 14.70 | 11.18 | 12.51 | 51.24 | 6.28 |
|  | *Morus notabilis* | 173.48 | 422,824 | 34.49 | 17.88 | 16.62 | 11.56 | 61.85 | 6.16 |
|  | *Prunus mume* | 208.95 | 355,740 | 41.61 | 17.77 | 23.84 | 18.21 | 56.16 | 12.11 |
|  | *Prunus persica* | 210.13 | 235,610 | 29.77 | 17.98 | 11.78 | 13.13 | 54.14 | 6.09 |
|  | *Pyrus bretschneideri* | 321.12 | 416,659 | 56.11 | 27.31 | 28.81 | 11.00 | 54.50 | 6.26 |
|  | *Ziziphus jujuba* | 284.69 | 632,000 | 52.46 | 22.34 | 30.12 | 12.32 | 61.91 | 7.73 |

**Table S3.** Summary of regulatory elements and interactions inferred by FunTFBS

| Group | Species | TF number (with binding motifs) | Regulatory element number (genome-wide) | Regulatory element number (in promoter) | TF number (in network) | Target gene number (in network) | Regulation number |
| --- | --- | --- | --- | --- | --- | --- | --- |
| BOP clade | *Brachypodium distachyon* | 236 | 144,761 | 17,589 | 235 | 7,044 | 16,108 |
|  | *Brachypodium stacei* | 230 | 143,220 | 18,072 | 230 | 6,709 | 16,325 |
|  | *Hordeum vulgare* | 191 | 198,060 | 5,992 | 188 | 2,857 | 5,472 |
|  | *Leersia perrieri* | 227 | 170,826 | 33,415 | 226 | 9,262 | 29,731 |
|  | *Oryza brachyantha* | 202 | 111,481 | 25,988 | 202 | 8,637 | 23,043 |
|  | *Oryza glaberrima* | 230 | 220,225 | 58,231 | 230 | 18,158 | 55,766 |
|  | *Oryza longistaminata* | 137 | 69,443 | 9,056 | 135 | 4,577 | 7,669 |
|  | *Oryza sativa subsp. japonica* | 247 | 299,203 | 50,021 | 245 | 13,784 | 43,224 |
|  | *Phyllostachys heterocycla* | 216 | 497,257 | 9,121 | 216 | 4,051 | 7,979 |
| Brassicales | *Aethionema arabicum* | 432 | 252,689 | 46,172 | 429 | 8,946 | 40,907 |
|  | *Arabidopsis halleri* | 460 | 150,913 | 31,635 | 454 | 7,461 | 28,860 |
|  | *Arabidopsis lyrata* | 562 | 364,983 | 69,748 | 561 | 13,560 | 64,675 |
|  | *Arabidopsis thaliana* | 598 | 327,942 | 66,414 | 596 | 12,567 | 64,185 |
|  | *Arabis alpina* | 394 | 324,092 | 40,440 | 392 | 10,081 | 37,116 |
|  | *Boechera stricta* | 562 | 421,352 | 64,973 | 560 | 11,732 | 59,714 |
|  | *Brassica napus* | 587 | 1,345,483 | 225,267 | 587 | 40,672 | 199,454 |
|  | *Brassica oleracea* | 545 | 633,268 | 76,044 | 545 | 17,973 | 70,491 |
|  | *Brassica rapa* | 560 | 575,852 | 84,317 | 559 | 18,083 | 78,596 |
|  | *Camelina sativa* | 635 | 1,444,148 | 217,768 | 635 | 36,680 | 199,087 |
|  | *Capsella grandiflora* | 541 | 191,046 | 40,312 | 539 | 8,988 | 38,597 |
|  | *Capsella rubella* | 564 | 301,464 | 55,862 | 563 | 11,143 | 51,755 |
|  | *Carica papaya* | 267 | 68,937 | 12,039 | 263 | 4,139 | 9,385 |
|  | *Eutrema salsugineum* | 544 | 415,850 | 68,065 | 543 | 12,334 | 60,815 |
|  | *Raphanus sativus* | 533 | 474,756 | 65,452 | 533 | 15,646 | 62,395 |
|  | *Sisymbrium irio* | 523 | 235,396 | 50,970 | 522 | 12,470 | 47,715 |
|  | *Tarenaya hassleriana* | 421 | 329,792 | 48,413 | 420 | 10,816 | 43,374 |
|  | *Thellungiella parvula* | 562 | 274,729 | 31,543 | 553 | 6,187 | 29,089 |
| Fabales | *Arachis duranensis* | 260 | 544,668 | 36,834 | 260 | 9,586 | 27,159 |
|  | *Arachis ipaensis* | 262 | 770,086 | 44,481 | 262 | 10,749 | 32,461 |
|  | *Cajanus cajan* | 300 | 281,050 | 33,131 | 300 | 9,179 | 26,802 |
|  | *Cicer arietinum* | 317 | 476,894 | 29,965 | 316 | 7,516 | 25,291 |
|  | *Glycine max* | 360 | 1,481,756 | 70,062 | 360 | 16,752 | 58,219 |
|  | *Glycine soja* | 325 | 620,622 | 51,841 | 325 | 13,943 | 41,372 |
|  | *Lotus japonicus* | 296 | 326,050 | 46,988 | 295 | 14,041 | 38,627 |
|  | *Medicago truncatula* | 308 | 282,088 | 29,835 | 308 | 10,022 | 26,151 |
|  | *Phaseolus vulgaris* | 327 | 612,096 | 28,143 | 327 | 7,519 | 24,032 |
|  | *Trifolium pratense* | 292 | 231,882 | 18,086 | 291 | 5,899 | 15,542 |
|  | *Vigna angularis* | 270 | 263,881 | 26,873 | 268 | 6,895 | 21,274 |
|  | *Vigna radiata* | 270 | 305,604 | 24,005 | 269 | 6,121 | 19,210 |
| Lamiales | *Dorcoceras hygrometricum* | 257 | 665,443 | 22,715 | 254 | 6,854 | 18,283 |
|  | *Mimulus guttatus* | 285 | 402,993 | 28,647 | 280 | 6,824 | 22,199 |
|  | *Sesamum indicum* | 305 | 168,320 | 14,457 | 301 | 4,163 | 11,267 |
|  | *Utricularia gibba* | 238 | 45,099 | 10,865 | 212 | 3,439 | 9,416 |
| Malpighiales | *Linum usitatissimum* | 299 | 139,254 | 26,916 | 297 | 8,378 | 21,459 |
|  | *Manihot esculenta* | 337 | 373,381 | 26,359 | 336 | 7,634 | 21,591 |
|  | *Populus trichocarpa* | 365 | 308,983 | 30,200 | 364 | 9,260 | 24,538 |
|  | *Ricinus communis* | 294 | 124,787 | 13,817 | 284 | 5,040 | 11,309 |
|  | *Salix purpurea* | 386 | 352,343 | 34,318 | 384 | 9,809 | 28,550 |
| PACMAD clade | *Dichanthelium oligosanthes* | 225 | 191,267 | 18,809 | 225 | 6,984 | 17,135 |
|  | *Eragrostis tef* | 191 | 141,771 | 12,193 | 191 | 6,167 | 11,081 |
|  | *Oropetium thomaeum* | 176 | 60,036 | 7,886 | 175 | 4,789 | 7,089 |
|  | *Panicum virgatum* | 280 | 582,055 | 54,759 | 280 | 20,604 | 51,495 |
|  | *Setaria italica* | 241 | 153,610 | 15,922 | 239 | 6,979 | 15,135 |
|  | *Setaria viridis* | 240 | 166,064 | 16,595 | 240 | 7,020 | 15,638 |
|  | *Sorghum bicolor* | 236 | 161,447 | 20,979 | 236 | 6,995 | 19,440 |
|  | *Zea mays* | 229 | 167,926 | 14,239 | 229 | 6,242 | 12,921 |
|  | *Zoysia pacifica* | 220 | 180,012 | 34,656 | 219 | 12,878 | 33,452 |
| Rosales | *Fragaria vesca* | 279 | 82,406 | 14,006 | 274 | 5,178 | 12,209 |
|  | *Morus notabilis* | 303 | 229,269 | 33,716 | 302 | 8,196 | 22,976 |
|  | *Prunus mume* | 323 | 190,470 | 20,660 | 316 | 6,940 | 16,283 |
|  | *Prunus persica* | 319 | 183,278 | 19,400 | 316 | 6,640 | 15,416 |
|  | *Pyrus bretschneideri* | 335 | 351,962 | 10,286 | 313 | 2,741 | 7,265 |
|  | *Ziziphus jujuba* | 338 | 391,480 | 28,014 | 337 | 7,824 | 20,748 |

**Table S4.** The resources released in this work

| Module | Description | Content | URL |
| --- | --- | --- | --- |
| *cis*-Map | A genome browser set up for users to browse genome alignments, conservation data and regulatory elements for 63 plants. | Pairwise genome alignments  Multiple genome alignments  Conserved elements  PhastCons scores  PhyloP scores  Conserved TFBS  FunTFBS | <http://plantregmap.cbi.pku.edu.cn/cis-map.php> |
| Network | A portal for users to retrieve and visualize regulations. | Regulations inferred by conserved TFBS and FunTFBS | <http://plantregmap.cbi.pku.edu.cn/network.php> |
| Tool  (FunTFBS) | A stand-alone package for users to screen for functional TFBS using the FunTFBS method. | Basic version | <http://plantregmap.cbi.pku.edu.cn/funtfbs.php> |
|  |  | Development version | <https://github.com/gao-lab/FunTFBS.git> |
| Download | A portal for users to download genome alignments, conservation data, regulatory elements and regulations. | Download individual file via HTTP | <http://plantregmap.cbi.pku.edu.cn/download.php> |
|  |  | Batch download via FTP | <ftp://ftp.cbi.pku.edu.cn/pub/database/PlantRegMap/> |
